# Two-photon imaging induces brain heating and calcium microdomain hyperactivity in cortical astrocytes

**DOI:** 10.1101/321091

**Authors:** Elke Schmidt, Martin Oheim

**Author notes:** corresponding author, −21 (lab). CLASSIFICATION: Major category: Biological Sciences Minor categories: Neuroscience / Physiology. AUTHOR CONTRIBUTIONS: ES and MO designed and performed research, ES analyzed the data, MO wrote the paper.

## Abstract

Unraveling how neural networks process and represent sensory information and how this cellular dynamics instructs behavioral output is a main goal in current neuroscience. Two-photon activation of optogenetic actuators and fluorescence calcium (Ca^2+^) imaging with genetically encoded Ca^2+^ indicators allow, respectively, the all-optical stimulation and readout of activity from genetically identified cell populations. However, these techniques expose the brain to high near-infrared light doses raising the concern of light-induced adverse effects on the biological phenomena being studied. Combing Ca^2+^ imaging of GCaMP6f-expressing cortical astrocytes as a sensitive readout for photodamage and an unbiased machine-based event detection, we demonstrate the subtle build-up of aberrant microdomain Ca^2+^ signals in fine astroglial processes. Illumination conditions routinely being used in biological two-photon microscopy (920-nm excitation, 100-fs regime, ten mW average power) increased the frequency of microdomain Ca^2+^ events, but left their amplitude, area and duration rather unchanged. This increase in local Ca^2+^ activity was followed by Ca^2+^ transients in the otherwise silent soma. Ca^2+^ hyperactivity occurred without overt morphological damage. Surprisingly, at the same average power, continuous-wave 920-nm illumination was as damaging as fs pulses, indicating a linear, heating-mediated (rather than a highly non-linear) damage mechanism. In an astrocyte-specific IP_3_-receptor knock-out mouse (IP_3_R2-KO), Near-infrared light-induced Ca^2+^ microdomains signals persisted in the small processes, underpinning their resemblance to physiological IP_3_R2-independent Ca^2+^ signals, while somatic activity was abolished. Contrary to what has generally been believed in the field, shorter pulses and lower average power are advantageous to alleviate photodamage and allow for longer useful recording windows.

**SIGNIFICANCE STATEMENT:** Imaging the fine structure and function of the brain has become possible with two-photon microscopy that uses ultrashort-pulsed infrared laser light for better tissue penetration. The high peak energy of these light pulses has raised concerns about photodamage resulting from multi-photon processes. Here, we show that the time-averaged rather than the peak laser power matters. At wavelengths and with laser powers now commonly used in neuroscience brain damage occurs as a consequence of direct infrared light absorption, i.e., heating. To counteract brain heating we explore a strategy that uses even shorter, more energetic pulses but a lower time-averaged laser power to produce the same image quality while making two-photon microscopy less invasive.

## INTRODUCTION

Two-photon (2P) excitation fluorescence microscopy (1) is the method of choice for imaging structural and functional dynamics in the intact brain (2-4). Addressing the role of genetically identified cell populations in a circuit-specific manner has become possible through the advent and continuous improvement of genetically encoded calcium (Ca^2+^) indicators (GECIs) (5-7). Considerable efforts are being made to enhance the speed, spatial resolution and field-of-view of 2P Ca^2+^ imaging, but for a given GECI the user has to trade off gains in one of these parameters against losses in others, unless the number of excitation spots and/or the average laser power 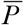 per spot are increased.

2P fluorescence excitation requires high instantaneous peak intensities, which has prompted concerns about non-linear photobleaching (8) and photodamage resulting from excited-state absorption restricted to the high-intensity focus (9, 10). Several studies have established a steep intensity threshold for this non-linear damage. The occurrence of aberrant, light-induced Ca^2+^ signals provides a sensitive readout for assessing damage (9–11). Average laser powers of the order of tens of mW per excitation spot are generally being considered safe (11), and up to 250 mW have been reported in multi-spot excitation schemes (12). How different pulse parameters (pulse length, inter-pulse interval, peak power) affect this damage threshold has remained controversial.

In the current work, we demonstrate an unexpected photodamage that affects physiological astrocytic Ca^2+^ signals, occurs upon 920-nm excitation at low mW powers, and involves a linear power dependence. Unbiased automated analysis of Ca^2+^ signals revealed a subtle but measurable build-up of spontaneous, microdomain Ca^2+^ activity in the fine processes of GCaMP6f-expressing cortical astrocytes under conditions routinely used for 2P brain imaging. Near-infrared (NIR) illumination increased the frequency whilst only slightly affecting the amplitude and cellular volume fraction encompassed by these Ca^2+^ transients. This frequency increase occurred without overt morphological damage. The build-up of aberrant microdomain Ca^2+^ signals was followed by Ca^2+^ activity in the otherwise silent soma. Pausing acquisitions reduced light-induced Ca^2+^ hyperactivity, indicating its dependence on the total light dose rather the timing with respect to the first image. Longer, less energetic pulses and higher average power (resulting in the same signal) were equally harmful. As pulse stretching did not decrease the photoinduced Ca^2+^ signals, as would have been expected for a non-linear damage mechanism, we postulated a linear (heating-mediated) damage process. Consistent with a one-photon absorption process, continuous-wave 920-nm illumination was equally damaging to small astrocyte processes as were fs pulses.

Our experiments demonstrate that one- rather than two-photon NIR-absorption is limiting in the now popular wavelength band above 900 nm. This has important consequences for imaging fluorescent proteins and functional indicators as well as for studies using optical stimulation for optogenetic activation. Thus, a careful optimization of the pulse length whilst reducing the average laser power better preserves the biology under study.

## RESULTS

### ‘Spontaneous’ astrocyte Ca^2+^ signals increase during two-photon recordings

In the barrel cortex of adult mice (P33-P46) sparsely expressing GCaMP6f (13) in a subset of astrocytes (**Fig. 1***A* and **Fig. S1**), 2-photon imaging at 920 nm revealed a plethora of spontaneous, asynchronous and spatially confined Ca^2+^ signals (**Fig. 1** and **Supplementary Movie S1**). These microdomain Ca^2+^ signals occurred in the absence of neuronal stimulation. They were astrocyte-intrinsic because they persisted when blocking neuronal action potentials with TTX (1 µM) and they were also observed in pure astrocyte culture (data not shown).

We acquired time-lapse image series (at 0.5 Hz) from a single equatorial plane encompassing the soma at diffraction-limited spatial resolution (146 nm/px). Regions of elevated Ca^2+^ (**Fig 1***A*, *bottom*) and their corresponding Ca^2+^ transient waveforms (**Fig. 1***B*, *top*) were extracted by an unbiased machine-based detection procedure, see **Fig. S2** and **Online methods**. The ‘summed activity’ trace (showing the cumulative activity in the processes) was dominated by a large number of small Ca^2+^ transients and a few large events and was otherwise flat, as expected for a predominantly localized, asynchronous Ca^2+^ activity. Soma showed little, if any, spontaneous activity during the first 4 min of recording. Ca^2+^ signals were rare (11.25 ± 1.76 events/min), short-lived (1.90 ± 0.10 s duration), of small amplitude (Δ*F*/*F*_0_ = 3.02 ± 0.12) and they were confined to tiny sub-regions of individual astrocyte branches (mean area, 9.26 ± 0.52 µm^2^, vs. 3969 ± 302 µm^2^ total astrocyte area, i.e., ∽0.2% of the surface area; median ± SEM for 8 cells during 4 min recording). Our results are similar to those reported by others (14-17).

**Fig. 1.**
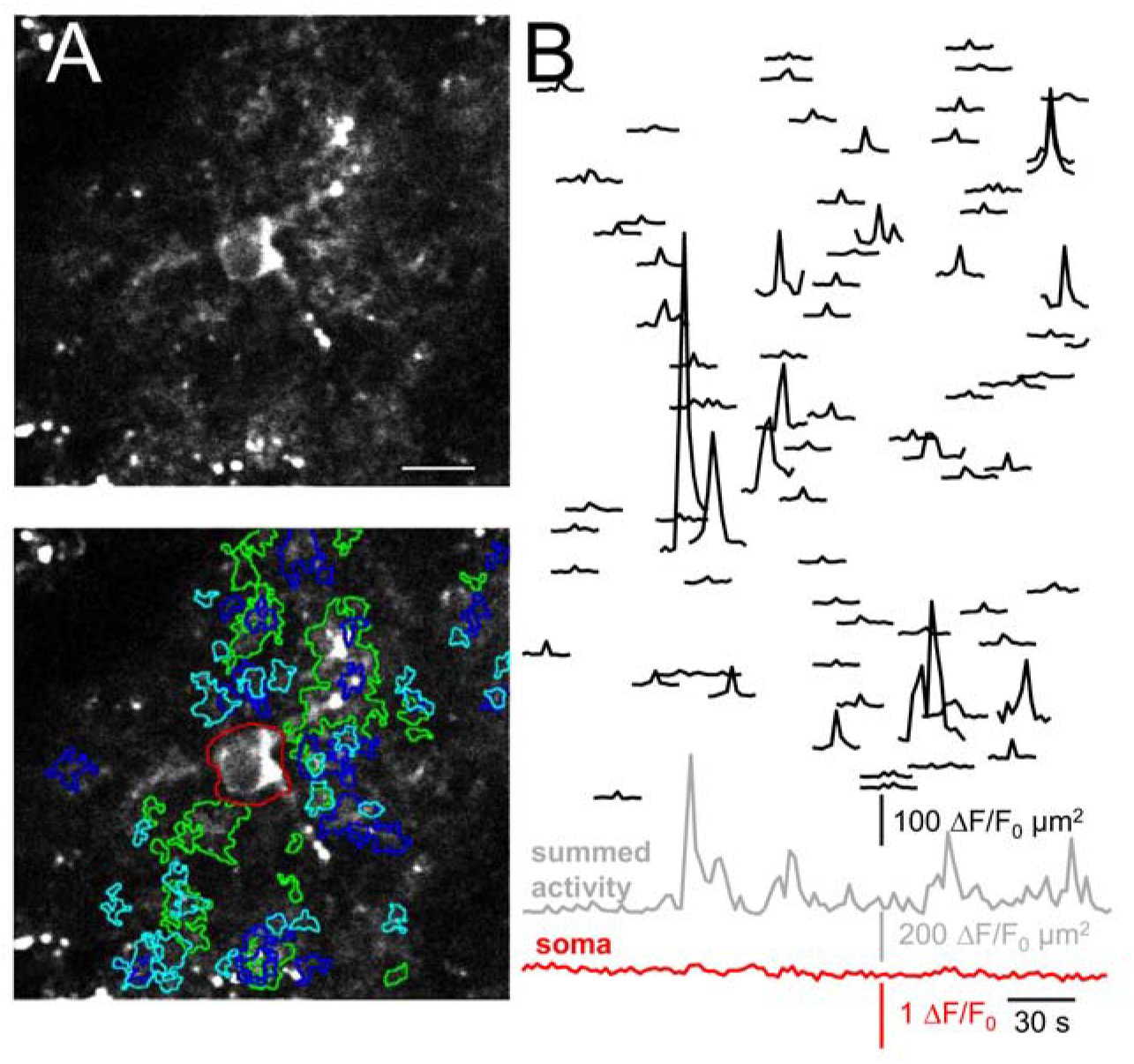
Properties of spontaneous astrocyte Ca^2+^ signals. (*A*) Diffraction-limited 920-nm excited two-photon (2P) fluorescence image across a single equatorial plane of a GCaMP6f-expressing astrocyte in an acute slice of the murine barrel cortex. Imaging depth 83 µm. *Top*, morphological overview from the time-average of 120 consecutive frames. *Bottom*, regions of interest (ROIs) corresponding to the detected Ca^2+^-transients (color for better discrimination). Overlapping ROIs represent activity occurring in the same area at different time points. *Red* central ROI is the soma. Scale bar, 10 µm. (See Movie S1). (*B*) *Black*, area-scaled fluorescence traces of spontaneous Ca^2+^ events (Δ*F*/*F*_0_ multiplied by the area of the corresponding ROI) ordered, from *top* to *bottom*, according to their distance to soma. See Fig. S2 and Online Methods for details on the detection algorithm. *Bottom*, *grey*, summed activity over all events in the astrocyte processes. *Red* trace somatic activity, the soma was silent throughout the recording.

Raster plots on which we ordered active regions according to their distance-to-soma showed Ca^2+^ transients throughout the neuropil with no obvious localization preference. When we extended the recording duration beyond the initial 4 min, we recognized a first subtle and then increasingly pronounced Ca^2+^ hyperactivity in the fine processes, **Fig. 2***A*, *top*. This hyperactivity is easily detected as a rise of the ‘summed activity’ trace that measures the cumulative activity within the processes (**Fig. 2***A*, *bottom*, grey trace). Somatic Ca^2+^ hyperactivity was observed secondary to the photodamage triggered in the cell periphery, (**Fig. 2***B*, *bottom*, red trace). With increasing recording time, astrocytic microdomain Ca^2+^ signals occurred repeatedly within the same subcellular domains, **Fig. 2***C*. Microdomain events remained initially relatively stereotyped in terms of their area, peak amplitude and duration but their frequency almost doubled (×1.87) when comparing p1 (0–4 min) and p2 (4–8 min). Following 16 min of continuous recording, microdomain Ca^2+^ transients in the processes tripled in frequency and had a 40%-increased d*F*/*F*_0_ amplitude, compared to the beginning (p1), **Fig. 2***D*. Other parameters remained unaltered, indicating that photodamage merely triggered more events. We conclude that both event frequency (×2.94) and summed activity (×7.5) of the processes are *bona fide* markers for light-induced microdomain Ca^2+^ hyperactivity. On the macroscopic level, longer recordings produced a net increase in somatic Ca^2+^ activity (×14.5, p3 vs. p1), **Fig. 3***E* and **Supplementary Movie S2**. Aberrant Ca^2+^ signals occurred in the absence of visible morphological damage, and mice expressing EGFP rather than GCaMP6f in an astrocyte-specific manner showed no obvious photoactivation (data not shown). Ca^2+^ hyperactivity was observed in astrocytes expressing GCaMP3 (18), a GECI displaying a higher basal fluorescence but reduced Δ*F*/*F*_0_ amplitude (19), excluding GCaMP6f-specific effects (data not shown).

The Ca^2+^ signal obtained when lumping together the entire neuropil to one single ROI mirrored the evolution with time of the summed activity of all individual detected events in the cell periphery (**Fig.S3**). Thus, (*i*), although highly branched and segregated into a tortious and diffusionally distant volume, the fine processes behave as a single compartment as far as damage is concerned, and, (*ii*), the soma behaves differently. The simple ‘center-vs-surround’ analysis hence provides a facile readout of aberrant light-induced Ca^2+^ activity in the soma and processes, respectively.

Taken together, our results demonstrate the vulnerability of small astrocyte processes to the exposure with pulsed near-infrared (NIR) light. Imaging conditions routinely used for biological 2P microscopy trigger astroglial Ca^2+^ hyperactivity detectable in both processes and soma.

**Fig. 2.**
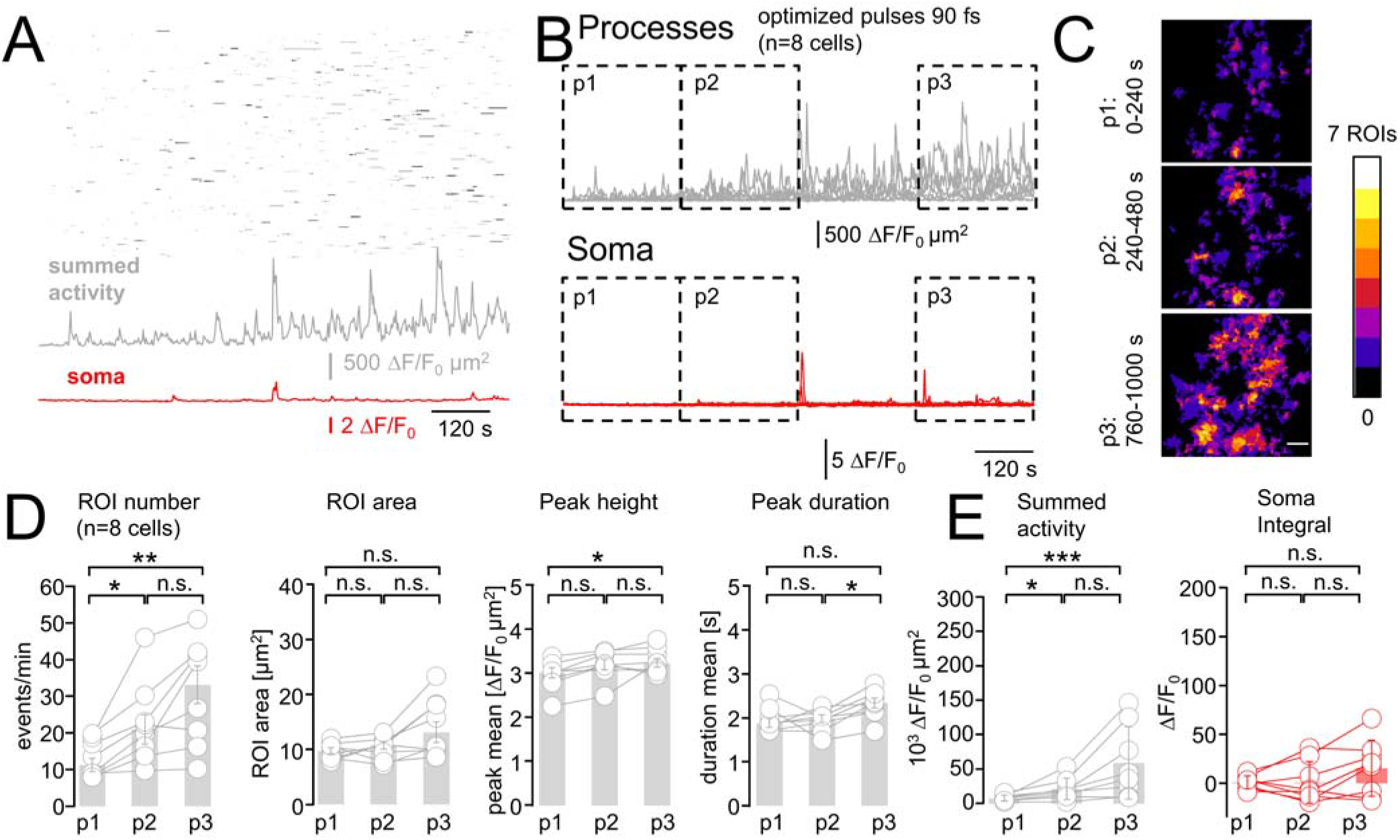
2P excitation at 920 nm triggers a subtle but detectable Ca^2+^ hyperactivity. (*A*) *Black*, raster plot of the dat in Fig. 1*B*, along with its further evlution during a recording lasting more than 16 min (500 frames, 0.5 Hz). *Grey*, increase of the summed activity over all processes. *Red*, the initially silent soma becomes progressively active. (*B*) Summed peripheral (*grey*) and somatic (*red*) activities under the same conditions as in (A) for *n* = 8 cells in *N* = 6 mice. Mean activity was quantified during periods p1: 0–240 s, p2: 240–480 s, p3: 760–1000 s, *boxed*. (C) Pseudo-color overlay of the events detected during p1, p2 and p3 for the cell in (A). Ca^2+^-activity increasingly occurs at similar spatial locations during p2 and p3. Scale bar, 10 µm. (*D*) Parameters characterizing Ca^2+^ transients during p1, p2 and p3. Bar plots show population median ± SEM, symbols graph individual cells. Event frequency increases from p1 to p2 and between p1 and p3 (two-sided nonparametric Wilcoxon-Mann-Whitney two-sample rank test). Light-induced Ca^2+^ signals resemble spontaneous Ca^2+^ microdomains, which makes them difficult to distinguish. (*E*) *Grey*, comparison of the summed activity in the astrocyte processes, i.e., integral of the traces shown in (B), reveals a net increase between p1 vs. p2 and p1 vs. p3. *Red*, the somatic activity, corresponding to the red traces in (B), remains significantly unchanged during all recording periods, although a trend to progressively higher activity is seen. *: *P* < 0.05, **: *P* < 0.01, ***: *P* < 0.005, n.s.: not significant.

### Pulse stretching does not reduce photodamage

For a given fluorophore, microscope and signal, we can reduce photodamage by modifying either the pulse frequency *f* or duration τ and adjusting the average laser power 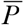 accordingly (i.e., keeping either 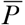 *f* = *const*. or 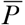/t = *const*., respectively). Longer pulses will reduce the pulse energy but require higher 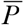. However, whether longer pulses are *per se* better for biological two-photon imaging has been a matter of debate. The outcome depends on whether photobleaching and photodamage increase more or less rapidly with 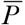 than the signal. Longer pulses will be neutral if damage processes scale with a power-exponent *m* (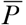^*m*^) of two, like 2P-excited fluorescence (9) whereas they will be beneficial if higher-order photodamage (*m* > 2) dominates (10). Conversely, longer pulses having a lower peak power and require a higher average power will exacerbate damage processes having a *m* < 2, e.g., when one-photon absorbers are present (20) or if tissue heating occurs (21).

We directly compared Ca^2+^ transients upon 920-nm excitation with 90- (as before) and 172-fs pulses. 90-fs pulses were obtained by a careful optimization of the excitation optical path and an optimal pre-chirping in a wavelength- and objective-dependent manner so as to attain shortest pulses in the sample plane (see **Methods**). 172-fs pulses correspond to the standard ‘midline’ manufacturer tuning of the DeepSee^TM^ pulse compressor at 920 nm.

To our surprise, longer pulses and higher laser power (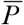 was adjusted so that astrocytes had an equal GCaMP6f signal for either pulse length, **Fig.S4**) did not reduce photodamage, **Fig. 3***A*. Aberrant microdomain Ca^2+^ signals were readily detectable throughout the astrocyte processes, **Fig. 3***B*, and somatic activation prevailed, **Fig. 3***D*. Quantifying the activity during initial (p1) and terminal 4-min segments (p3) revealed by and large identical results for 172- and 90-fs pulses in the processes (cumulative activity increased ×6.0 vs. ×7.5, p1 vs. p3), **Fig. 3***C*. The overall somatic activity in p3 was not significantly different between pulse lengths, **Fig. 3***E*, although somatic Ca^2+^ transients arrived earlier and were of larger amplitude with longer pulses. Our results clearly exclude highly non-linear photodamage as a mechanism for triggering aberrant astrocyte Ca^2+^ activity.

**Fig. 3.**
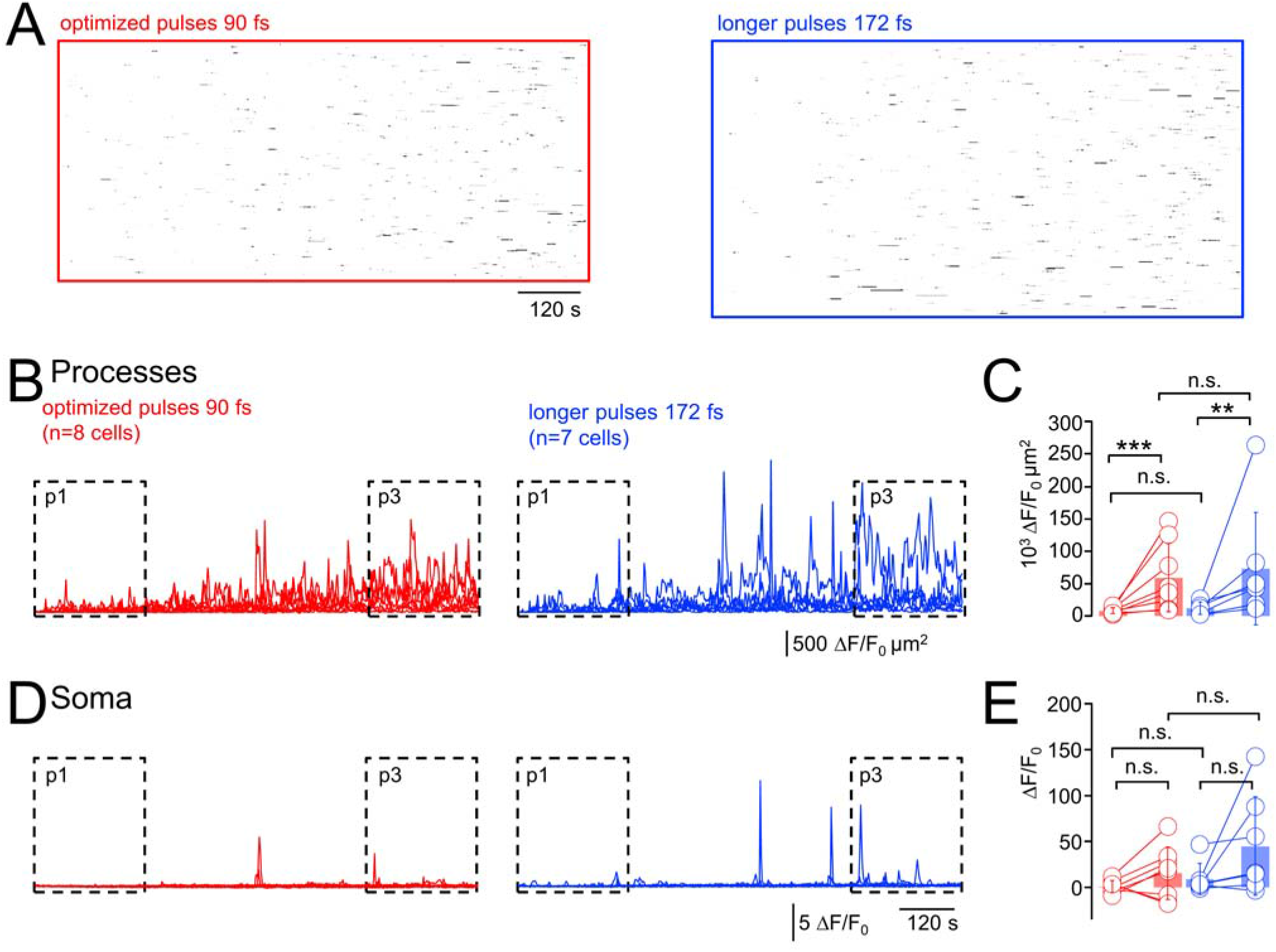
Longer, less energetic pulses do not reduce light-evoked Ca^2+^-hyperactivity. (*A*) *Right*, raster plot of 920-nm light-induced Ca^2+^ hyperactivity as in Fig. 2*A*, but for a different cell. Pulses were carefully optimized as in the both previous figures to achieve the shortest possible pulses under the objective (see Online methods), resulting in 90-fs pulse length (sech^2^). See Movie S2 for a recording of this cell. *Left*, same for a cell excited with 172-fs pulses, i.e., with lower peak power but higher average laser power to maintain a constant initial signal, i.e., 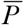^2^/τ = *const*. (*B*) *Red*, summed activity as in Fig. 2*B* (*n*=8 cells/6 mice at 90 fs) and side-to-side comparison with summed activity upon 172-fs excitation (*blue*). (*C*) Ca^2+^ activities in p3 compared to p1 (defined as in Fig. 2*B*) increase (median ± SEM) for both 90-fs (*red*, same data as in Fig. 2*E*) and 172-fs pulses (*blue*, *n* = 7 cells/6 mice). Furthermore, activity was undistinguishable between short and long pulses for p1 (validating our experimental paradigm) but also during p3, i.e., contrary to what was expected for a highly non-linear photodamage no significant difference was found for different pulse energies at (one-sided non-parametric Wilcoxon-Mann-Whitney two-sample rank test). (*D*) Somatic Δ*F*/*F*_0_ traces for 90-fs (*red*, corresponding to Fig. 2*B*) and 172-fs pulses (*blue*). (E), Statistical analysis showed that the somatic activity was indistinguishable between 90-fs (*red*, corresponding to Fig. 2 E) and 172-fs pulses (*blue*), both during p1 and p3. *: *P* < 0.05, **: *P* < 0.01, ***: *P* < 0.005, n.s.: not significant.

### CW-illumination at 920-nm is as damaging as fs-pulses to the fine processes

How can we explain that longer pulses and higher peak powers are not more damaging? We hypothesized that contrary to common belief, in the 100-fs and 920-nm excitation regime, photodamage is *not* dominated by highly non-linear processes like excited-state absorption, absorption from the triplet state, or indirect effects that occur via ROS production through mitochondrial two- and three-photon absorption or lipid oxidation, but rather through direct NIR absorption. This is plausible, because both water and lipid absorption increase by more than one order of magnitude between 800 and 900 nm (**Fig.S5**), and so does the risk of focal heating (*m* = 1) (21, 22). Thus, different from the short wavelengths (700–800 nm) (23) that traditionally have been used for 2P excitation of small chemical indicators, the longer wavelengths now commonly used for exciting fluorescent proteins and optogenetic activators might well result in significant absorption and heat production. A simple, testable hypothesis for a linear damage process is that depositing the same average power 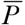 with a continuous-wave (CW) 920-nm laser (i.e., no pulsing) should be equally damaging as pulsed illumination with 2P fluorescence excitation. This is what we tested next, after having ascertained the effectiveness of our de-modelocking procedure (see **Online methods** and **Fig.S6***A*, *B*).

We designed a “pump-probe” experiment, in which we first imaged cells during 2 min with 90-fs pulses. We then pursued during 10 min with either pulsed or continuous-wave (CW) illumination (same 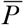). As no fluorescence is excited with CW illumination, we read out the resulting Ca^2+^ signals during another 2-min ‘probe’ period with pulsed excitation. In a third variant (negative control), we shuttered the laser during 10 min to allow the cells to recover, **Fig. 4***A*. For each cell, we graphed raster (**Fig. 4***B*) and cumulative-activity plots (**Fig. 4***C, E*). As before (c.f., **Fig. 2, Fig. 3**), ongoing fs-pulsed illumination led to synchronized ‘microdomain Ca^2+^ activity, **Fig. 4***D* as well as activation of somatic Ca^2+^ signals, **Fig. 4***F*. As hypothesized, CW illumination produced an amount of peripheral Ca^2+^ hyperactivity that was indistinguishable from pulsed excitation, **Figs. 4***D*, *F*, *grey* traces. Shuttering the laser between acquisitions virtually abolished this light-induced activity, indicating that the total light dose, not a once triggered irreversible damage cascade produced the aberrant Ca^2+^ signals, **Figs. 4***D*, *F*, *black* traces. Interestingly, and different from the astrocyte processes, CW illumination did not produce measurable somatic hyperactivity, **Fig. 4***F*. Together, our data provide compelling evidence for laser-induced heating being a source of peripheral astrocyte Ca^2+^ hyperactivity.

**Fig. 4.**
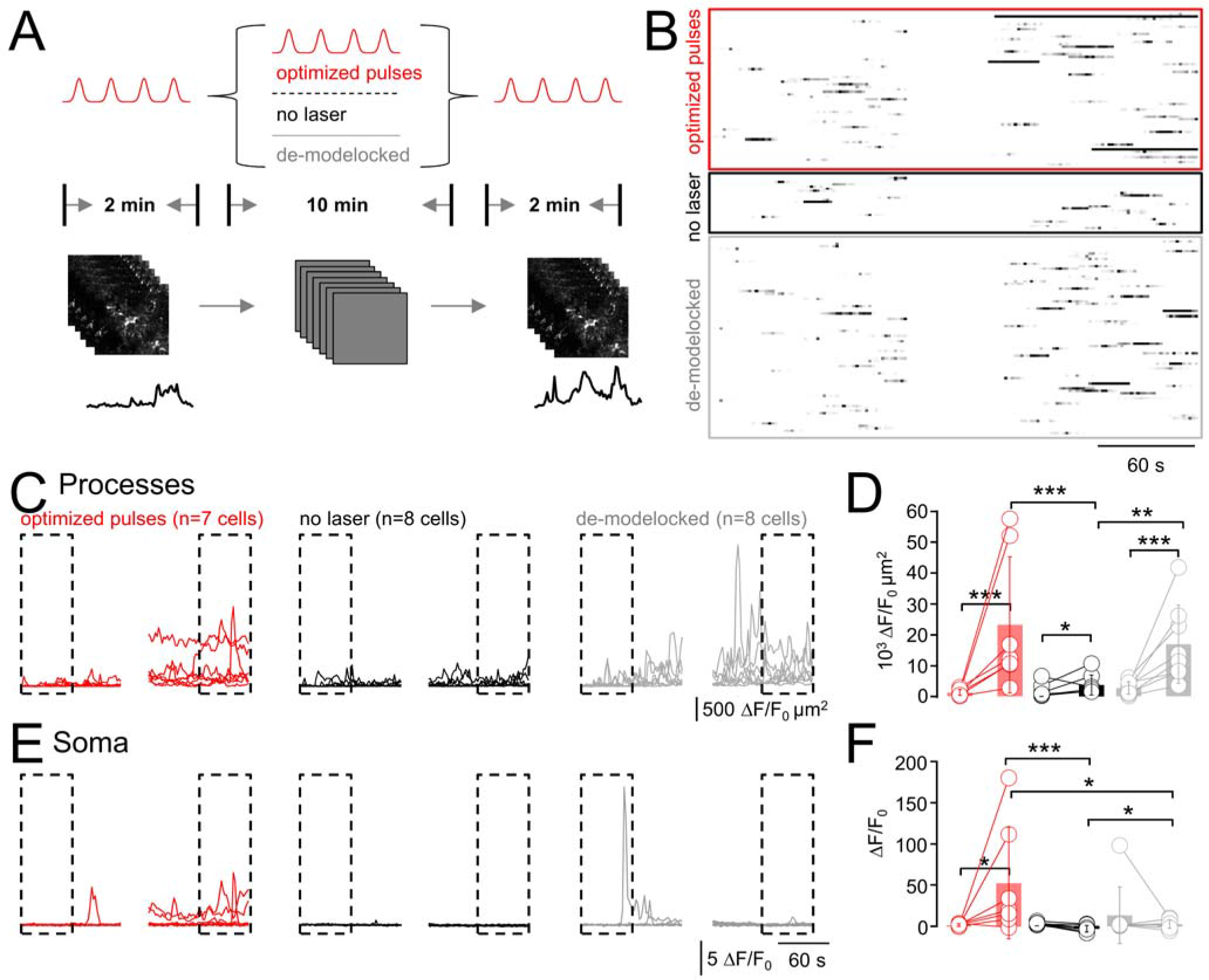
Light-induced Ca^2+^ hyperactivity depends on total light dose, not the pulse energy. (A) Schematic representation of the experiment design. Spontaneous Ca^2+^-transients were recorded during a 2-min control period (p1; 920 nm, 90 fs, 0.5 Hz). Then, during 10 minutes, three different protocols were applied: (*i*), the cells were either imaged as in the preceding figures (i.e., same scenario as before) or, (*ii*), the laser was shuttered and the cells allowed to recover from the previous recording. In a 3^rd^ variant, (*iii*), we de-modelocked the laser to expose the cells to continuous-wave (CW) radiation with the same average power 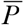 as during pulsed excitation (see Online Methods and Fig. S6). The respective impact of these different protocols on the astrocytic Ca^2+^ activity was read out in another 2-min recording (same parameters as p1). (*B*) Raster plots illustrate the outcome of the three different protocols. (C) Cumulative Ca^2+^ activity traces in the astrocyte processes during p1 and p3. (D) Statistical analysis of data shown in (C). Activity increases for all conditions between the beginning of p1 (0–60s) and end of p3 (660–720 s; one-sided non-parametric Wilcoxon-Mann-Whitney two-sample rank test). Of note, the Ca^2+^-activity in the processes during p3 is indistinguishable for exposure to pulsed vs. CW light, both of which are significantly higher than with the laser shuttered. All experiments started from initial activity levels during p1. (*n.s*. not shown for clarity). (E), Somatic Δ*F*/*F*_0_ traces for the three scenarios and statistical analysis in (F). For CW illumination, somatic activity is significantly higher in p3 compared to either no or pulsed excitation during p2, respectively. *: *P* < 0.05, **: *P* < 0.01, ***: *P* < 0.005, n.s.: not significant.

### Somatic but not peripheral photodamage is IP_3_-receptor mediated

What are possible damage mechanisms in the processes and soma? Perhaps reactive oxygen species (ROS) affect Ca^2+^ handling by oxidation of the inositol 3-phosphate receptor (IP_3_R) and sensitization of endoplasmic reticulum (ER) Ca^2+^ release to promote perisomatic Ca^2+^ oscillations, mitochondrial Ca^2+^ uptake (24) and release (25). As the IP_3_R2 is the major, if not only, IP_3_R expressed in astrocytes (26), we took advantage of an IP_3_R2-knock-out (KO) mouse (27) to test if IP_3_R sensitization was involved in mediating somatic damage. As spontaneous Ca^2+^ signals persist in the processes in IP_3_R2-KO mice (28) we speculated if imaging-evoked somatic and peripheral Ca^2+^ signals could be distinguished through their IP_3_R2-dependence. As before, we ascertained for all experiments equal 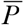(**Fig.S6***C*) and mean GCaMP6f-fluorescence at the beginning of the recordings (**Fig.S6***D*).

Time-lapse 2P imaging of spontaneous Ca^2+^ activity GCaMP6f-expressing astrocytes in slices of IP_3_R2-KO mice produced heat-induced microdomain Ca^2+^ transients in the cell periphery as in the WT, **Fig. 5***A*, while the somatic Ca^2+^ hyperactivity was abolished, **Fig. 5***C*. Analysis of initial and terminal 1-min segments respectively, revealed a significant heat-induced Ca^2+^ activity in the processes, *blue* on **Fig. 5***B*, both with respect to the beginning of the same recording and when compared to the second period in wild-type mice when shuttering the laser (data reproduced from **Fig. 4*D***). However, the same robust activity increase in the processes did not lead somatic responses in the KO, as if an IP_3_-mediated integration of the Ca^2+^ hyperactivity over different astrocyte branches was required to produce aberrant somatic signals, or ROS production caused by non-linear photodamage causes IP_3-_dependent somatic Ca^2+^ hyperactivity which is mechanistically different from the heat-triggered Ca^2+^ increase in the processes.

**Fig. 5.**
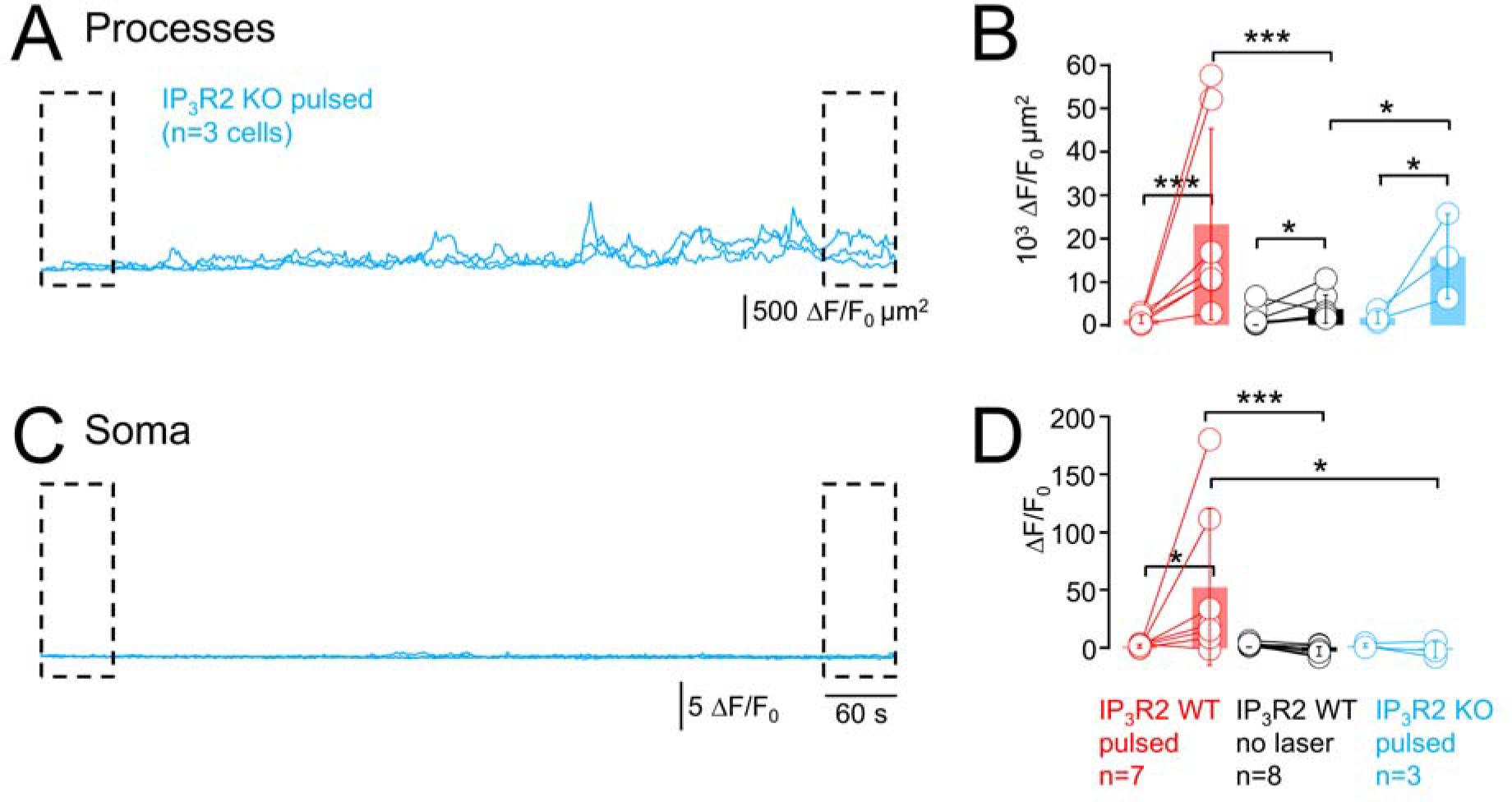
Somatic but not peripheral Ca^2+^-hyperactivity is IP_3_R2 receptor dependent. (A) Cumulative Ca^2+^ activity in the astrocyte processes during a 12-min recording (920 nm, 90 fs, 0.5 Hz) of GCaMP6f-expressing astrocytes from IP_3_R2 KO mice shows a pronounced light-induced Ca^2+^-hyperactivity in the processes (*n* = 3 cells from 2 mice). (B) Comparison of cumulative activity in the processes of IP_3_R2 KO mice during beginning of p1 (0–60s) and end of p3 (660–720s) (p1 and p3 same as in in Fig. 4, *light blue* graphs, one-sided non-parametric Wilcoxon-Mann-Whitney two-sample rank test). No significant difference is observed for either p1 or p3 between IP_3_R2 KO or wild-type (WT) mice (data from Fig. 4*D*), whereas the KO data during p3 is significantly different from the same period when the laser was shuttered (data from Fig. 4*D*), confirming that peripheral Ca^2+^ hyperactivity persisted in the absence of the IP_3_-mediated signaling pathways. In contrast, (C), Soma of IP_3_R2 KO astrocytes are silent during the whole recording period in all cells. (D) In fact, somatic activity in IP_3_R2 KO mice is more similar to the WT astrocytes that were only imaged during p1 and p3 with the laser shuttered in between than to those that were exposed to continuous 2PEF imaging (c.f., Fig. 4*F*). *: *P* < 0.05, **: *P* < 0.01, ***: *P* < 0.005, n.s.: not significant.

## DISCUSSION

We used state-of-the art methods to image and analyze microdomain Ca^2+^ signals in cortical astrocytes to show that, (*i*), 2P-imaging under conditions usually considered safe led to an increase in spontaneous Ca^2+^ activity; (*ii*), at constant signal (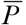^2^/t = const.), stretched pulses and increased average power failed to reduce photodamage, thus excluding non-linear damage mechanisms; (*iii*) 920-nm CW excitation evoked aberrant microdomain Ca^2+^ signals in the astrocyte processes with a comparable efficiency than pulsed excitation, confirming a one-photon absorption mechanism; (*v*), genetic ablation of the major astroglial IP_3_ receptor abolished somatic Ca^2+^ transients, but it did not affect heat-mediated damage in the fine processes, pointing to different damage mechanisms in the cell body and periphery.

### Brain heating as a damage mechanism

Heating was discussed as a possible damage mechanism in non-linear microscopies before the advent of the 2P microscope (29). Sheppard predicted permissive 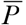 delivered to a single excitation spot around 35 mW and 100 mW for stationary illumination and fast scanning, respectively, which seems optimistic in view of reported biological damage thresholds closer to 10 mW (9-11). Comparing fs vs. ps pulses Schönle and Hell judged focal heating irrelevant at 750-800 nm (the dominant spectral window for exciting small-molecule chemical indicators), but neither they used a biological readout, nor is it clear if the biology under study would really tolerate the predicted 0.5-3°C-change (23). Morphological damage of unlabeled cultured cells exposed to fs and ps pulses (780 nm, 60 µs) followed an approximate 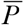^2^/τ-law, suggesting a damage exponent close to *m* = 2 (20), but more subtle forms of photodamage than visible destruction were not assessed in this study. For slightly longer wavelengths, Marcias-Romero *et al*. estimated a 1-5°C increase in the focus of a 1035-nm laser beam (80 MHz, 1.1 NA) in water for typical pixel dwell times <10 µs (30) and argued for wide-field illumination and low-repetition rate 2P imaging to reduce thermal damage. Using the temperature-dependent Stokes shift of Laurdan, a 1°C/100 mW change was reported for a CW 1064-nm trapping beam in the 20- to 200-mW regime (31). Along the same lines, Schmidt and co-workers predicted a temperature rise around 5°C/100 mW for various trapping beams based on a model taking into account light absorption in the neighborhood of the focus, outward heat flow and heat sinking by the glass surfaces of the sample chamber (32). A temperature increase of the same order of magnitude (1.8°C/100 mW) was measured with thermocouple probes and quantum-dot nanothermometers upon 2P single-spot scanning in the neocortex of awake or anesthetized mice, *in vivo*, and 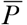 >250 mW produced irreversible thermal damage (21).

Our results underpin the importance of thermal damage for two-photon excitation in the biologically important spectral window around 920 nm and call for a necessary degree of caution when designing and interpreting experiments using 2P-brain imaging and photostimulation. Wavelengths >900 nm are now commonly being used for the excitation of fluorescent proteins, genetically encoded Ca^2+^ indicators (GECIs) and for the optogenetic activation of opsin-expressing cells. NIR light at these wavelengths is more invasive than often thought, particularly when imaging at high spatio-temporal resolution or when using multiple-spot schemes that increase the power deposit in tissue.

As thermal effects rely on the accumulation and dissipation of energy, they depend on 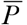, but also on parameters like the focal spot size, scanning parameters and the pulse frequency and shape. Moreover, the heat deposit will be additive in multi-spot, line-scanning, and holographic illumination schemes, and scanning with high spatial and temporal frequencies will intensify thermal damage. On the other hand, external factors like brain cooling due to a cranial window (33) or perfusion with temperature-controlled ASCF can contribute to equilibrating temperature gradients.

At a fixed pulse length, we expect lower pulse repetition rates, shorter pixel dwell times and re-scanning (i.e., temporal averaging on a µs time-scale) (34) to be beneficial. In view of thermal diffusion rates of ∽1 µm^2^/µs, scanning schemes in which neighboring pixels are excited in a spatially interspersed manner rather than sequentially will favor the dissipation of locally generated heat. A low overall duty cycle (i.e., recording from multiple, distant and small ROIs rather than from full frames, temporally spacing short bursts of images taken at high frequency (21)) should be beneficial to mitigate build-up of thermal damage. Random-access scanning (35-37) will be advantageous for applications where high spatio-temporal sampling is indispensible like imaging dendritic spines, astrocyte processes, or for 2P-STED microscopy.

Our work alerts the biological microscopist to the importance of other damage mechanisms that coexist with localized non-linear photodamage (10, 38) and it also calls for prudence in the ongoing quest for ever-longer wavelength IR-excited and red-shifted probes for deep tissue imaging, because water and lipid absorption (*µ*a) continue to increase. However, tissue scattering (*µ*s) is reduced at these wavelengths. Thus, the choice of the optimal fluorophore and excitation wavelength windows will be a trade-off between absorption and tissue scattering (39, 40). Finally, thermal damage is a particular concern for deep-tissue imaging because compensating for the exponential excitation losses with increasing depth will expose the tissue surface to very high laser powers that – even unfocused – will be harmful due to surface heat generation.

### Somatic Ca^2+^ signals are a sensitive readout for damage

Although somatic Ca^2+^ transients have been used as a proxy for astrocyte activity, ≥90% of spontaneous Ca^2+^ activity occurs in the cell periphery (14, 17, 41-43). We confirm this observation (Figs. 1, 2) and find that astrocyte somata are mostly silent under conditions of 2P-images that preserve physiological Ca^2+^ signaling. Even when neuronal action potentials were not blocked with TTX, somatic Ca^2+^ signals were sparse, making their appearance a sensitive and facile readout for damage, however, heat-induced alterations of microdomain Ca^2+^ activity in the cell periphery occurred earlier (Fig. 2) and were more difficult to spot.

Light-induced somatic Ca^2+^ transients in cortical astrocytes depend on IP_3_-mediated mechanisms (Fig. 5). In the cell periphery with its large membrane-to-cytosol ratio, other mechanisms prevail. Candidates are channels with thermosensitive gating in the physiological temperature range, like TRPV3 (44), STIM-1 (45), and a number of voltage-gated channels (46). Temperature variations around 37°C also sensitize TRPV4 channels that are expressed in roughly 1/3 of astrocytes (47) to diverse stimuli triggering channel opening (48). A different Ca^2+^-influx pathway potentially recruited following NIR illumination and regulated by oxidative stress is TRPM2 (49). Alternatively, heat-induced pore-formation or lipid rearrangements in the membranes of Ca^2+^ stores or the plasma membrane could mediate microdomain Ca^2+^ hyperactivity.

### Towards even shorter pulses?

Our study illustrates the potential benefit of shorter pulses and lower 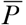 for 2P-excitation of GCaMP-based indicators at 920 nm and in the 100-fs regime. 90-fs were the minimum we could attain with our laser, microscope, objective and pre-compensation optics, but even shorter pulses might be beneficial (50, 51). This strategy will eventually be bounded by some shortest tolerable pulse length because the resulting high peak powers will favor non-linear photo-bleaching and -damage pathways. The demonstration that phase-optimized pulses with 28-fs length reduced the bleaching rate of EGFP by a factor of 4 while maintaining the same intensity of the fluorescence signal (52) let us predict that there might be room for a further reduction of NIR-induced damage for brain imaging with even shorter pulses.

## METHODS

Full experimental procedures are available online.

## ACKNOWLEDGEMENTS

We thank Cendra Agulhon (Paris) for providing the IP_3_R2-KO mouse line and immunofluorescence data, Frank Pfrieger (Strasbourg) for providing the GLASTcreER^T2^ mouse line, Marcel van ’t Hoff for programming the pulse-length adjustment in LABVIEW and Stéphane Dieudonné (Paris), Manfred Lindau (Göttingen/Cornell) and many other colleagues for helpful discussions. Patrice Jegouzo, Christophe Tourain (workshop) and the animal house team provided excellent technical assistance.

Financed by the European Union (FP6-STRP “AUTO-SCREEN”, FP7 ERA-NET NEURON “NANOSYN”, FP7 JPND “SYNSPREAD”, H2020 EUROSTARS “OASIS”), the FranceBioImaging large-scale national infrastructure initiative (FBI, ANR-10-INSB-04, *Investments for the future*) and the Region Ile de France (Cancéropôle, “EDISON”). The Oheim lab is a member of the *Ecole de Neurosciences de Paris* (ENP) and the C’nano Ile-de-France excellence clusters for neurobiology and nanobiophotonics, respectively.

This work contains supporting online material.

